# Integrating spatial, phylogenetic, and threat assessment data from frogs and lizards to identify areas for conservation priorities in Peninsular Malaysia

**DOI:** 10.1101/2021.04.07.438880

**Authors:** Kin Onn Chan, L. Lee Grismer

**Author notes:** Corresponding author’s.

## Abstract

Malaysia is recognized as a megadiverse country and biodiversity hotspot, which necessitates sufficient levels of habitat protection and effective conservation management. However, conservation planning in Malaysia has hitherto relied largely on species distribution data without taking into account the rich evolutionary history of taxa. This represents the first study that integrates spatial and evolutionary approaches to identify important centers of diversity, endemism, and bioregionalization that can be earmarked for conservation priorities in Peninsular Malaysia. Using georeferenced species occurrences, comprehensive phylogenies, and threat assessments of frogs and lizards, we employed a spatial phylogenetics framework that incorporates various diversity metrics including weighted endemism, phylogenetic diversity, phylogenetic endemism, and evolutionary distinctiveness and global endangerment. Ten areas of high conservation value were identified via the intersection of these metrics—northern Perlis, Langkawi Geopark, southern Bintang range, Cameron Highlands, Fraser’s Hill, Benom-Krau complex, Selangor-Genting complex, Endau-Rompin National Park, Seribuat Archipelago (Tioman and Pemanggil Islands), and southern Johor. Of these, Cameron Highlands requires the highest conservation priority based on severe environmental degradation, inadequately protected areas, and high numbers of endangered and evolutionary distinct species. Other areas, especially in the northwestern (states of Kedah and Penang) and northeastern regions (states of Kelantan) were not only identified as areas of high conservation value but also areas of biogeographic importance. Taken together, frogs and lizards demonstrate distinct east-west and north-south patterns of bioregionalization that are largely modulated by mountain ranges.

**Article Impact Statement:** The first study to use a spatial phylogenetic approach to identify areas for conservation priorities in Malaysia

## Introduction

Malaysia is a developing and megadiverse country located within the Western Sunda biodiversity hotspot (Myers et al. 2000; Mittermeier et al. 2005). Due to rapid economic growth in recent decades, forest cover in Peninsular Malaysia has been substantially reduced from as high as 80% in 1940 to 60% in 1971; at the end of 2016, only 44% of forest cover remained (Aiken 1994; FDPM 2016). To safeguard its natural and cultural resources, approximately 14% of the total land area has been designated as protected areas (PAs) and this figure is targeted to increase to 20% by 2025 in accordance with the National Policy on Biological Diversity (NRE 2016).

A major criterion for establishing PAs is the conservation of species and habitat based on scientific data (Hashim et al. 2017). Traditionally, conservation planning has predominantly relied on spatial patterns of biodiversity at the species level (Clements et al. 2008; Evans et al. 2018; Ratnayeke et al. 2018). However, these approaches ignore the rich evolutionary history of taxa and how they diversified across space and time. In contrast, instead of solely focusing on species distributions, spatial phylogenetic approaches also take into account the evolutionary relationships among organisms at all levels (Carnaval et al. 2014; Mishler et al. 2014; González-Orozco et al. 2015; Fenker et al. 2020). This integrative approach can provide a more holistic understanding of biodiversity patterns—for example, studies have shown that areas of high phylogenetic diversity or endemism are not necessarily areas of high species richness and endemism (Rosauer et al. 2009; Mishler et al. 2014). Therefore, conservation strategies that incorporate spatial data and evolutionary history can be better at ensuring the long-term persistence of entire lineages as opposed to individual species.

Numerous metrics have been developed to quantify and characterize biodiversity. Weighted endemism (WE) covers the study area with a grid of cells of equal size and weights the species richness of each cell by the inverse of its distribution range (Williams & Humphries 1994). Phylogenetic diversity (PD) is defined as the sum of branch lengths of a set of taxa in a phylogeny that serves as a proxy for the total amount of biodiversity for those taxa (Faith 1992). This measure also has the benefit of being robust to taxonomic change or uncertainty (Mace et al. 2003; Rosauer et al. 2009). Phylogenetic endemism (PE) combines WE and PD to provide a measure of endemism to phylogenetic diversity, i.e. the degree to which PD is spatially restricted (Rosauer et al. 2009; Carnaval et al. 2014). The relative contribution of a species to PD can also be used to represent evolutionary distinctiveness (ED), which reflects the degree of phylogenetic isolation/uniqueness for a particular species (Pavoine et al. 2005; Redding & Mooers 2006; Isaac et al. 2007). When ED is combined with extinction risk derived from the International Union for Conservation of Nature (IUCN) Red List, the Evolutionary Distinctiveness and Global Endangerment (EDGE) index can be used to prioritize species for conservation not only based on their likelihood of extinction, but also on their irreplaceability (Redding & Mooers 2006; Isaac et al. 2007; Daru et al. 2017, 2020). Each of these metrics is informative for a specific facet of biodiversity and when integrated within a unified framework across multiple taxonomic groups, can provide a more robust and comprehensive characterization of evolutionary history and biodiversity patterns that can be used to prioritize conservation initiatives (Posadas et al. 2001; González-Orozco et al. 2015).

Spatial phylogenetic approaches to conservation management have not been implemented before in Malaysia largely due to the lack of genetic resources in the past. However, over the last decade or so, surveys to unexplored as well as commonly explored areas and the increased use of genetic methods in biodiversity research have generated a wealth of spatial and phylogenetic data, especially for frogs and lizards (Chan & Grismer 2008; Grismer & Chan 2008; Chan & Ahmad 2009; Matsui 2009; Matsui et al. 2009, 2014; Chan et al. 2009, 2010a, 2010b, 2014, 2019, 2020; Grismer et al. 2010c, 2010a, 2011a, 2011b, 2014b, 2014a; Quah et al. 2011, 2017, 2019b, 2019a; Sumarli et al. 2015, 2016; Davis et al. 2016). These taxa have high extinction rates and relatively restricted ranges, which makes them ideal organisms for identifying areas of high conservation priority (Isaac et al. 2012; Barratt et al. 2017; Fenker et al. 2020; Gumbs et al. 2020). For this study, we examined patterns of phylogenetic diversity and endemism across a large swathe of frog and lizard species in Peninsular Malaysia (data for snakes were not included in the reptile dataset due to the lack of adequate spatial and genetic representation) to identify significant centers of biodiversity and patterns of bioregionalization that can inform conservation strategies.

## Methods

### Sampling

Spatial data in the form of point occurrences (GPS coordinates) were either obtained from literature or unpublished sources that have been vouchered by specimens or photographs. All points were curated, georeferenced, and verified by the authors to determine their accuracy. Species with no accompanying phylogenetic data were excluded. In total, our datasets included 470 georeferenced points from 70 species of frogs (63% of total diversity; Table S1) and 162 points from 85 species of lizards (47% of total diversity; Table S2). Global endangerment status was obtained from the IUCN red list database (IUCN 2019) and information on PAs was taken from the Malaysia Biodiversity Information System (https://www.mybis.gov.my/one/).

Single-locus mitochondrial phylogenies were constructed from genetic sequences obtained from GenBank (Tables S3, S4). The 16S rRNA gene was selected for frogs, while the NADH dehydrogenase 2 (ND2) gene was selected for lizards. These genes have the highest taxonomic coverage and are widely considered the most informative mitochondrial markers for their respective groups. A total of 607 and 408 sequences were obtained for frogs and lizards respectively, with each sequence representing a single species. Species and genera that are closely related to Peninsular Malaysian taxa but do not occur there were also included to avoid biases stemming from missing taxa. Sequences were aligned using MAFFT v7.471 (Katoh & Standley 2013) through the CIPRES Science Gateway (Miller et al. 2010) and subsequently checked for errors in Geneious v5.6 (Kearse et al. 2012).

### Analysis

The program IQ-TREE v2.0.5 (Minh et al. 2020) was used to perform model testing (Kalyaanamoorthy et al. 2017) and maximum likelihood phylogenetic estimation. Branch support was assessed with Ultrafast Bootstrapping (Hoang et al. 2017) via 1000 bootstrap replicates. The R package *phyloregion* (Daru et al. 2020) was used to calculate WE, PD, PE, and EDGE. Point data were converted to a community data frame at a resolution of 0.5 decimal degrees. To identify significant areas, we discarded areas that scored below the third quartile for each metric. We found this threshold to retain a reasonable and manageable number of areas—a second-quartile threshold retained too many areas for meaningful consideration. Retained areas for frog and lizard datasets were then combined across all metrics to identify areas that were common and disparate between both groups of taxa. Intersecting areas for frog and lizard datasets were considered areas of high conservation value.

The program Infomap Bioregions was used to identify bioregions and indicative species using an adaptive resolution method to account for incomplete species distribution data (Edler et al. 2017). Top indicative species are defined as species with the highest relative abundance, which is calculated as the ratio between the frequency of the species in the region and the frequency of the species in all regions (Edler et al. 2017). Because the Infomap analysis does not require a phylogeny, we used expanded occurrence datasets which include taxa that were previously excluded from the phyloregion analysis due to the lack of accompanying genetic material (Tables S5, S6). The Infomap analysis was implemented through the web application (https://bioregions.mapequation.org) with a maximum cell size of 0.5 degrees, number of trials = 1, and number of cluster cost = 1.00 as recommended by the developers.

## Results

Between 7–11 areas scored within the third quartile for both frog and lizard datasets across all metrics (Fig. 1). For frogs, three areas were highly significant as indicated by the overlap of all four metrics. For lizards, four areas were highly significant (Fig. 1). When frog and lizard datasets were combined, ten intersecting areas were identified as areas of highest conservation value (Fig. 2A). These include Langkawi Geopark (area 1; Fig. 2A), northern Perlis (area 2), southern Bintang range (area 3), Cameron Highlands (area 4), Fraser’s Hill (area 5), Benom-Krau complex (area 6), Selangor-Genting complex (area 7), Endau-Rompin National Park (area 8), Seribuat Archipelago (Tioman and Pemanggil Islands; area 9), and southern Johor (area 10).

**Fig. 1.**
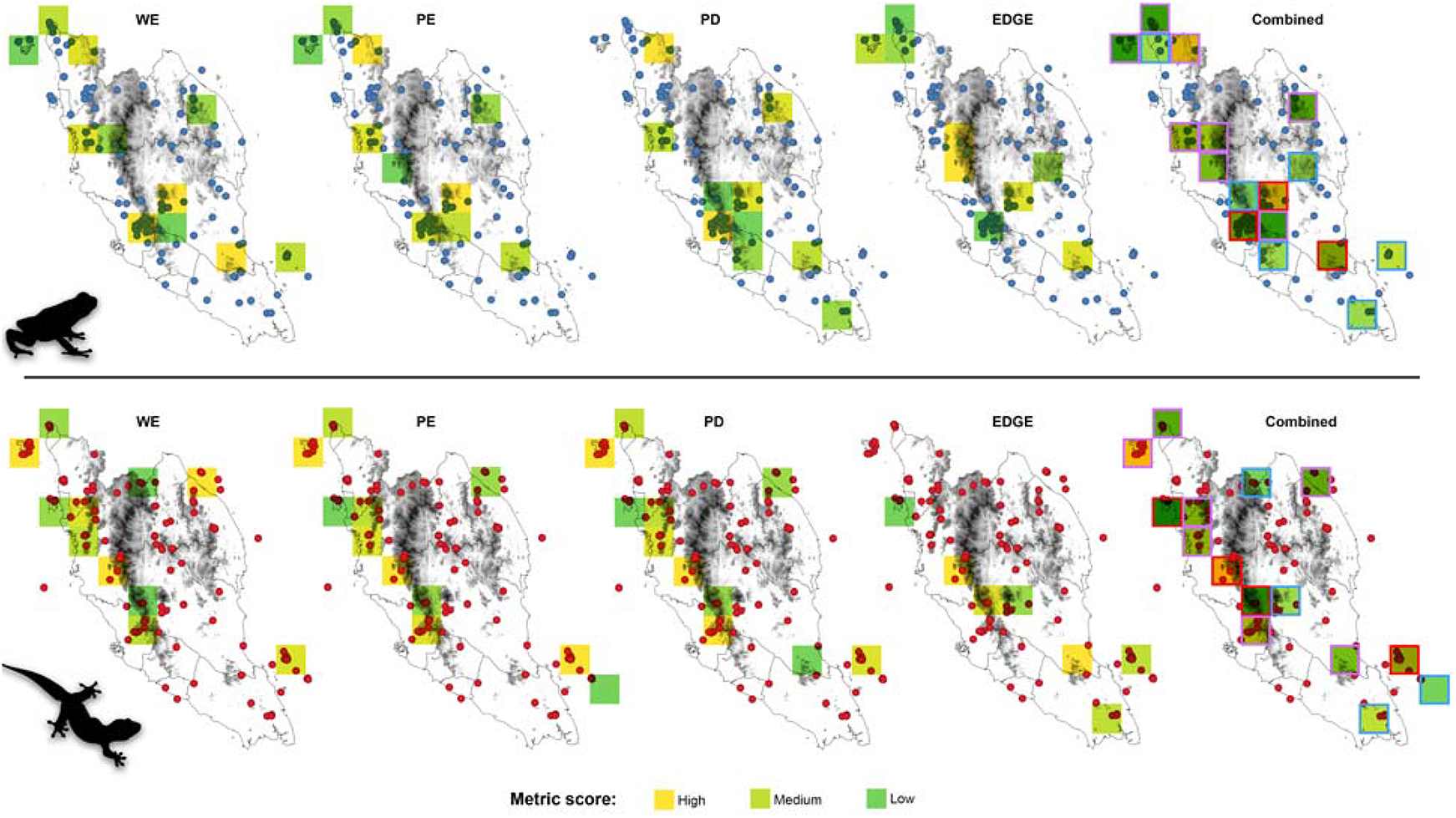
Distribution of areas that scored within the third quartile in weighted endemism (WE), phylogenetic diversity (PD), phylogenetic endemism (PE), and evolutionary distinctiveness and global endangerment (EDGE) for frogs (top) and lizards (bottom). Combined layers for all metrics (WE + PD + PE + EDGE) are in the last column. Cells outlined in red in the combined layers are areas supported by all four metrics; cells outlined in purple are supported by 2–3 metrics; cells outlined in blue are supported by only one metric.

**Fig. 2.**
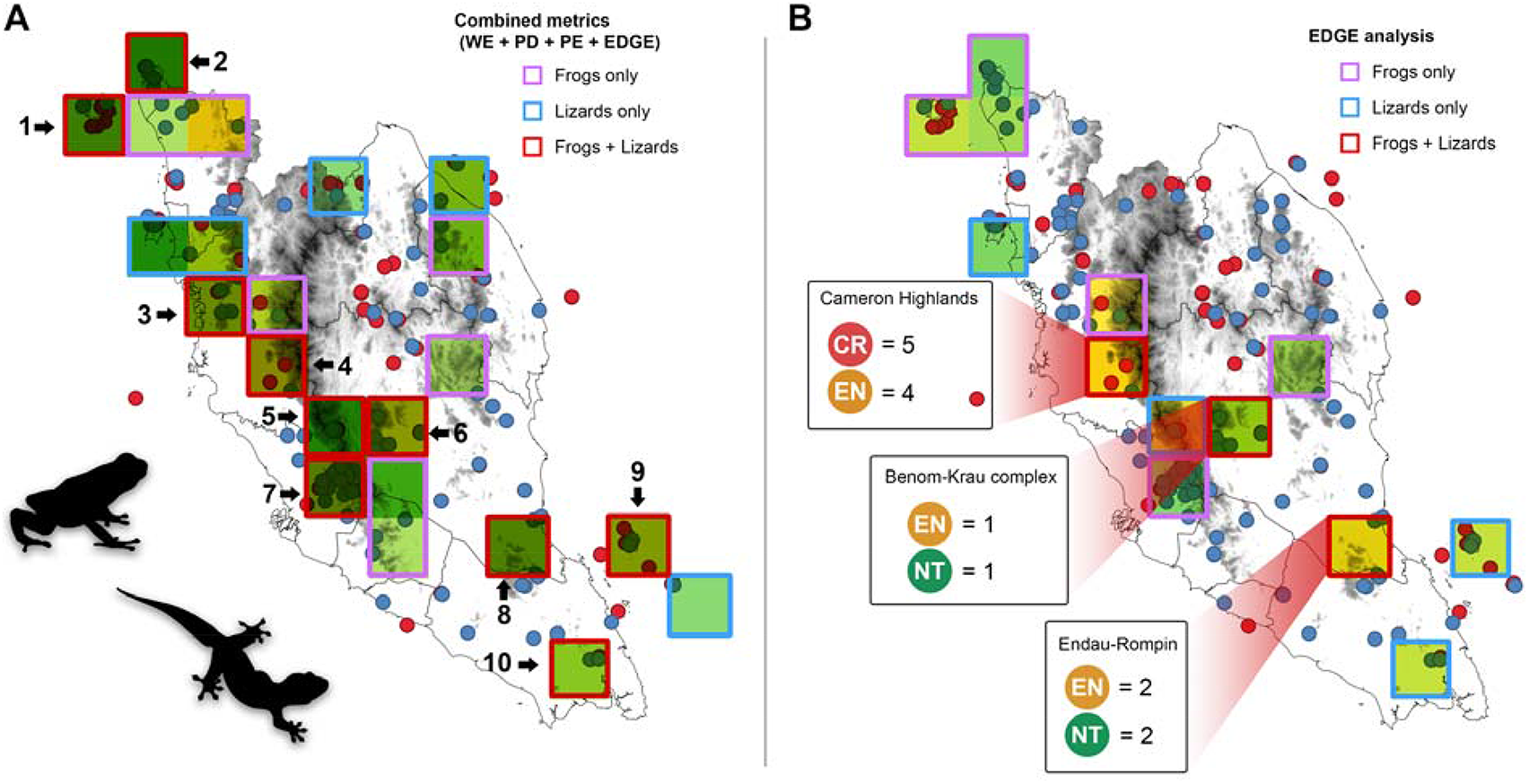
**A:** Areas of high conservation value that scored within the third quartile for all four metrics in both frogs and lizard datasets. Intersecting areas that are common to both datasets are outlined in red: Area 1 = Langkawi Geopark; 2 = northern Perlis; 3 = Southern Bintang range (Bukit Larut and Bintang Hijau); 4 = Cameron Highlands; 5 = Fraser’s Hill; 6 = Benom-Krau complex; 7 = Selangor-Genting complex; 8 = Endau Rompin National Park; 9 = Tioman and Pemanggil Islands; 10 = Southern Johor. Areas outlined in purple were only identified from the frog dataset; areas outline in blue were only identified from the lizard dataset **B:** Results of the EDGE-only analysis for the frog and lizard datasets combined. The number of near threatened (NT) and threatened (Endangered, EN; Critically Endangered, CR) species within each common area are shown. Blue circles = frog point occurrences; red triangles = reptile point occurrences.

Results from the EDGE-only analysis showed that among these areas, three were common between frogs and lizards: Cameron Highlands, Benom-Krau complex, and Endau-Rompin National Park (Fig. 2B). Cameron Highlands recorded the highest number of threatened species, followed by Endau-Rompin National Park, and Benom-Krau complex. The list of areas with the highest conservation value and their associated indicator species (inferred from the Infomap analysis), EDGE scores, PAs, and support metrics are presented in Table 1. Cameron Highlands has the highest number of threatened and evolutionary distinct species, exemplified by nine indicator species, all of which have high EDGE scores above 0.6 (Table 1). Benom-Krau complex (area 6), Selangor-Genting complex (area 7), Endau-Rompin National Park (area 8), and southern Johor (area 10) were supported by all metrics in the frog dataset, while Cameron Highlands (area 4), Fraser’s Hill (area 5), and Tioman-Pemanggil Islands (area 9) were supported by all metrics in the lizard dataset, indicating that these set of areas have particularly high conservation value for those respective groups. All ten areas are partially or fully protected by PAs of various categories (see Discussion for a more comprehensive review on these PAs).

**Table 1.**
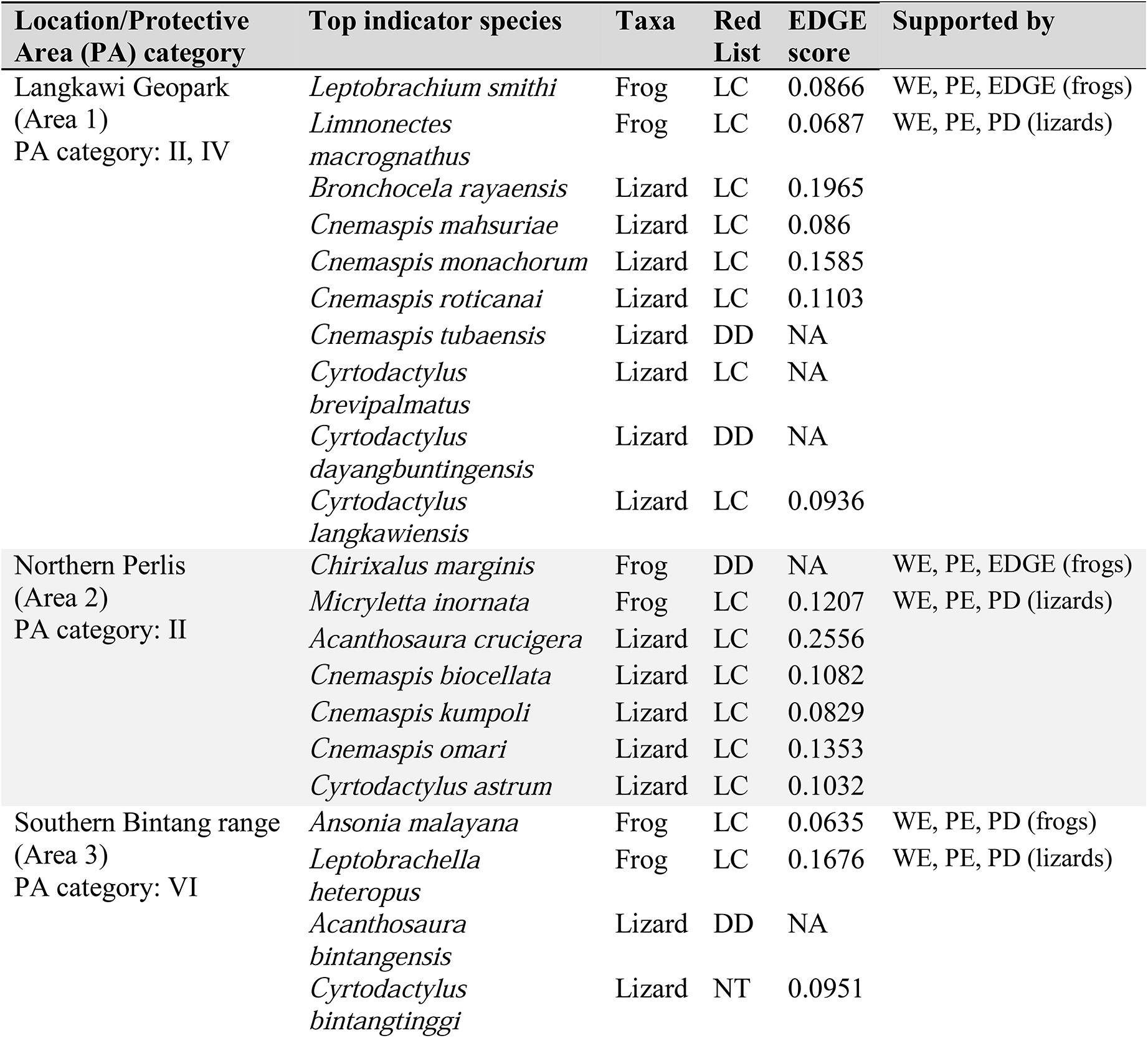
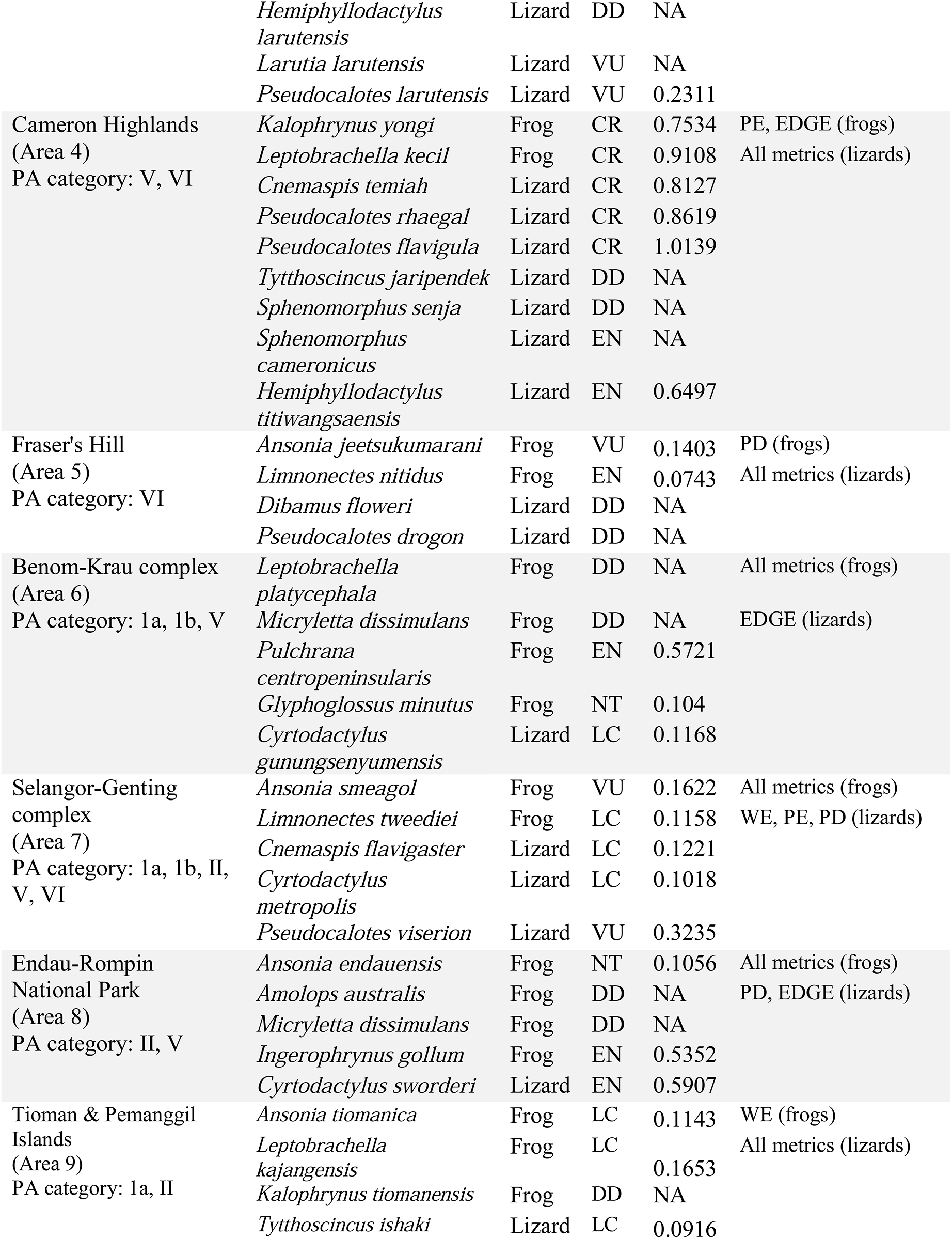
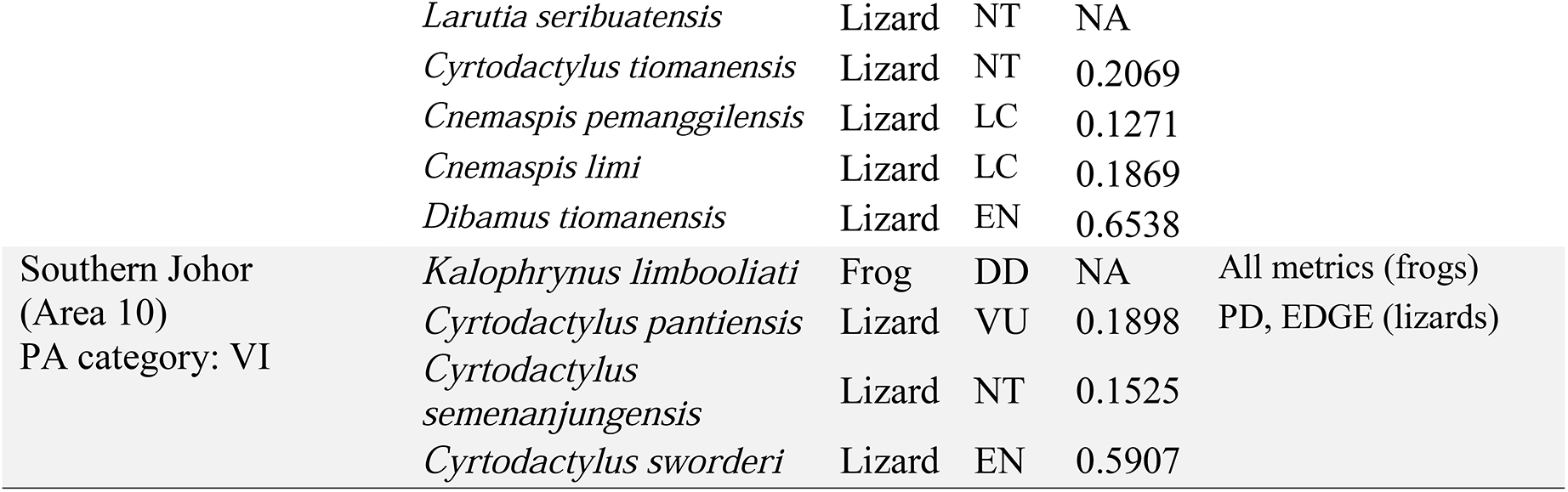
Areas of high conservation value identified from the combined frog and lizard datasets. The numbering of areas follows Figure 2. Top indicative species were inferred using the Infomap Bioregions program. CR = critically endangered, EN = endangered, VU = Vulnerable, NT = near threatened, LC = least concern, DD = data deficient, NA = data not available due to the lack of genetic or IUCN Red List data. Categories for protected areas follow IUCN: 1a = Strict Nature Reserve; 1b = Wilderness Area; II = National Park; III = Natural Monument or Feature; IV = Habitat/Species Management Area; V = Protected Landscape/Seascape; VI = Protected area with sustainable use of natural resources. WE = weighted endemism; PE = phylogenetic endemism; PD = phylogenetic diversity; EDGE = evolutionary distinctiveness and global endangerment.

The Infomap Bioregions analysis at cluster cost = 1 inferred seven regions for frogs (Fig. 3A). Several distinct regions were inferred: two areas in the northwestern region (areas 1 and 2; Fig. 3A), eastern region (areas 3 and 4), and southern region (areas 5 and 6). The inferred biogeographic regions for the lizard dataset at cluster cost = 1 were significantly more numerous (26 regions) and fragmented, indicating that lizard taxa are more range-restricted (Fig. 3B). To obtain a more conservative and comparable estimation, we performed a separate Infomap Bioregions analysis at a higher cluster cost of 2, which resulted in 16 regions (Fig. 3C). Similar to the frog dataset, the northwestern (area 7; Fig. 3C), eastern (areas 9–14), and southern regions (area 15) were also identified as distinct biogeographic regions. One additional region (area 8; Fig. 3C) was not identified from the frog dataset. A summary of significant biogeographic regions and their associated important areas and indicator species is presented in Table 2.

**Fig. 3.**
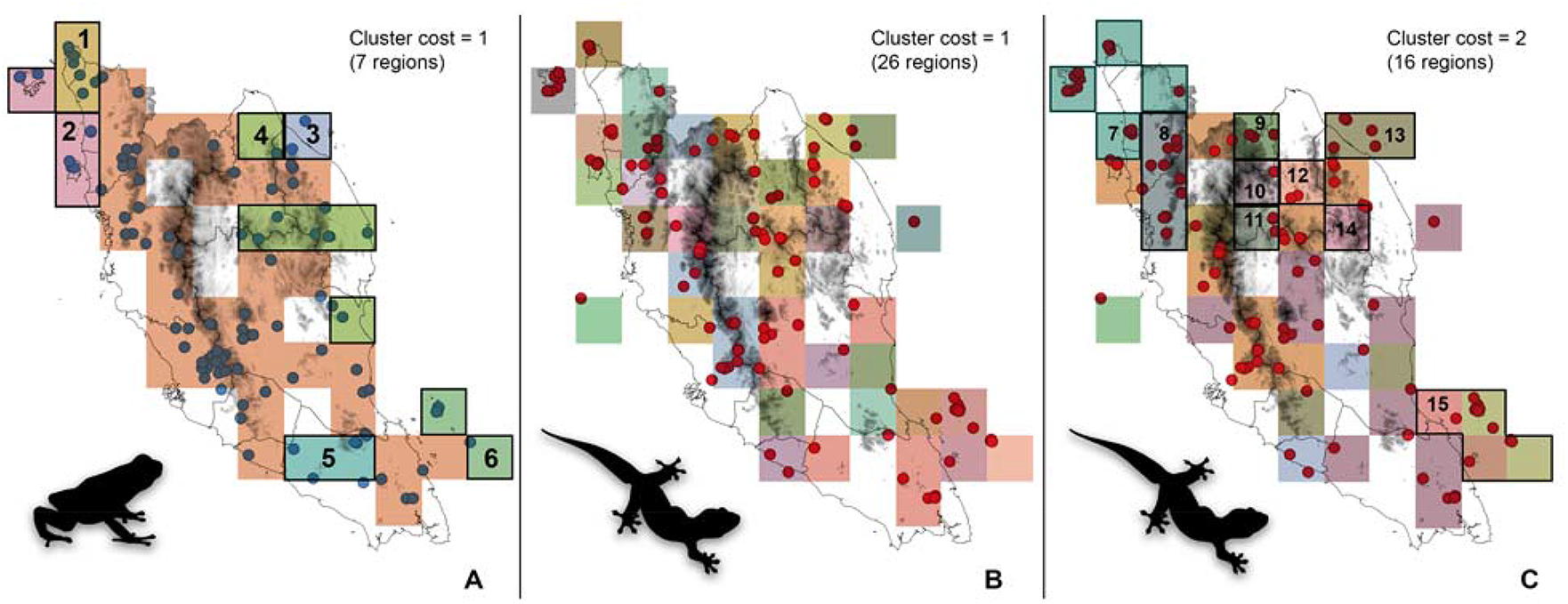
Important biogeographic regions inferred from the Infomap Bioregions analysis for frogs (A) and lizards (B, C) at differing cluster costs. Notable biogeographic regions for frogs are outlined in black and labeled 1–6, while notable regions for lizards are labeled 7–15. Blue circles = frog point occurrences; red triangles = reptile point occurrences.

**Table 2.**
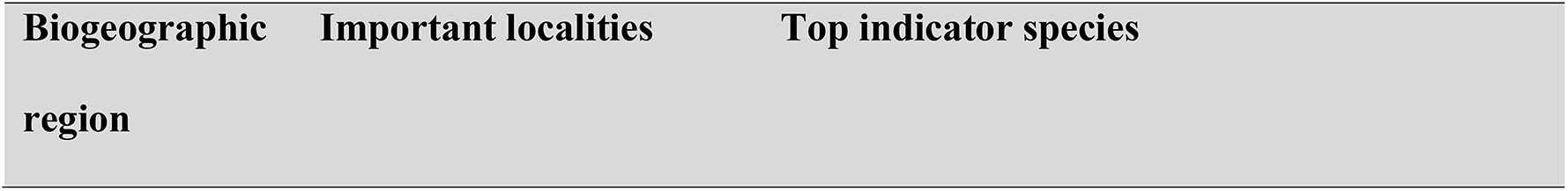
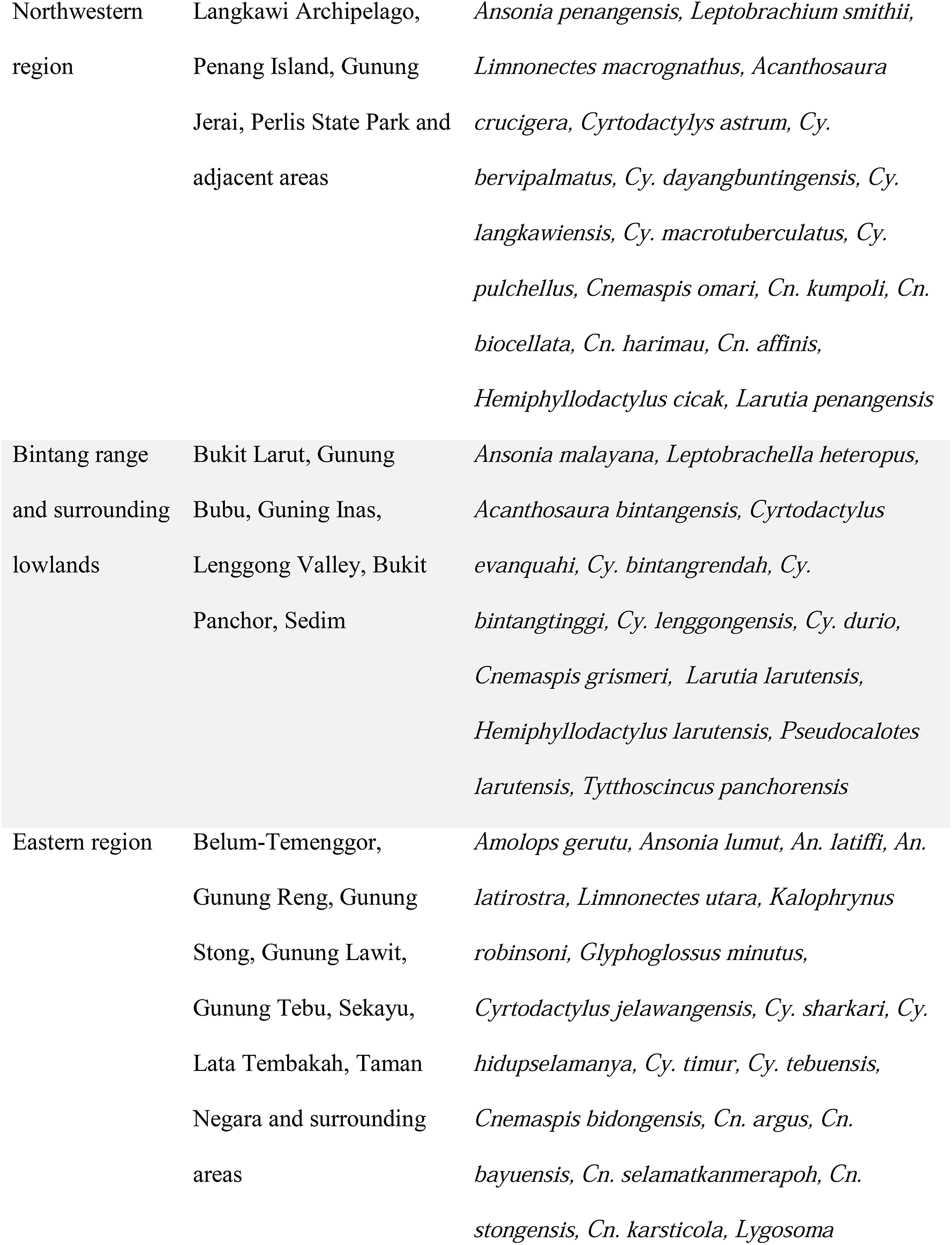
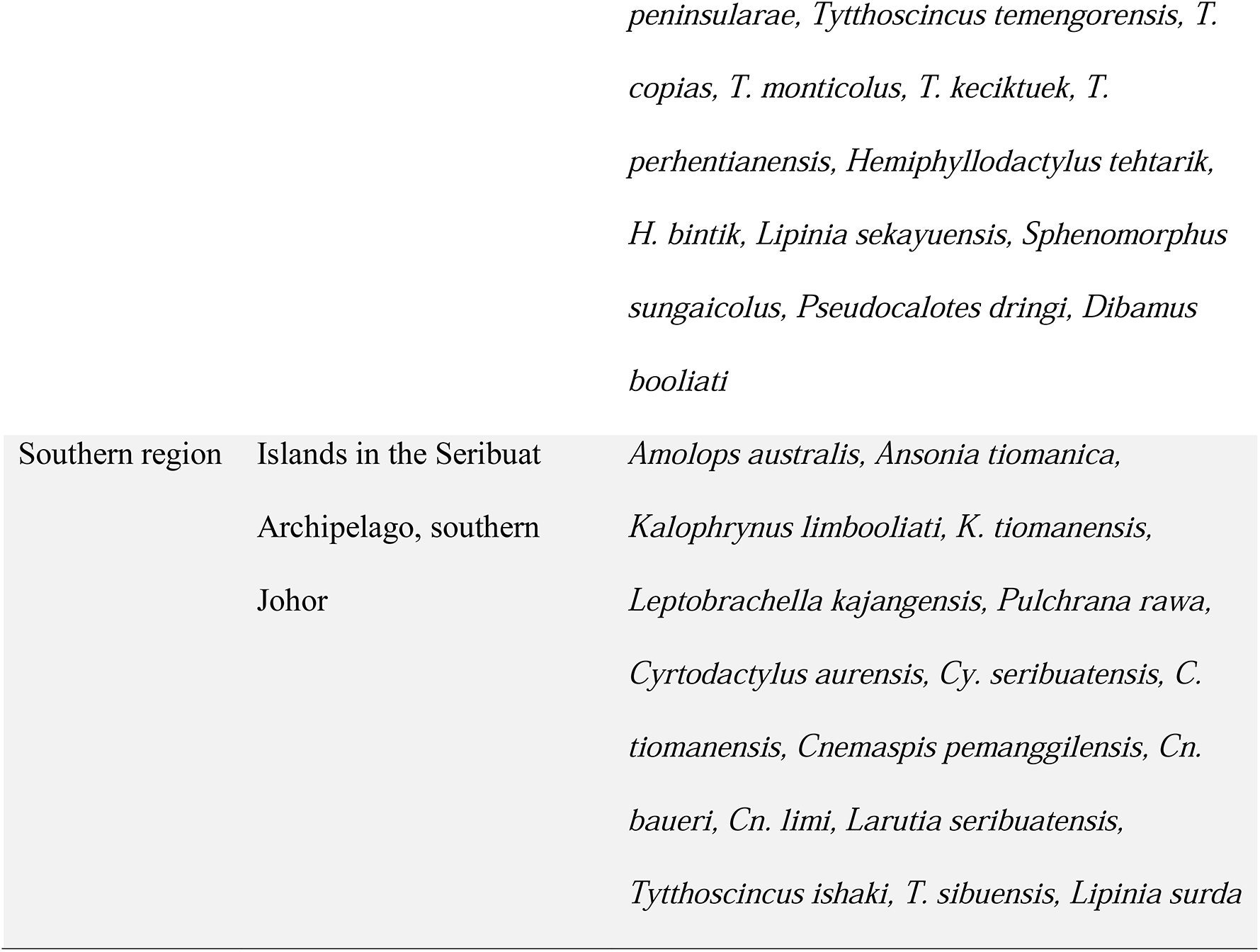
Significant biogeographic regions and their associated important areas and indicator species as inferred from the Infomap Bioregions analysis.

## Discussion

Although all ten areas of high conservation value (combined frog and lizard datasets) inferred from our data contained PAs (Fig. 2A; Table 1), the level of protection (PA category) for each area varied. The Benom-Krau complex (area 6; Fig. 2A), Endau-Rompin National Park (area 8), and Tioman-Pemanggil Islands (area 9) have the highest level of protection. Within the Benom-Krau complex, Gunung (=Mount) Benom and the Krau Wildlife Reserve are classified as a Strict Nature Reserve (category 1a) and Wilderness Area (category 1b), respectively (Table 1). Human visitation, use, and impacts are strictly limited in these areas. Endau-Rompin and Tioman Island are designated as a National and Marine Park, respectively (category II), which provides large-scale protection of the entire functional ecosystem. However, protection for area 9 excludes Pemanggil Island, which harbors numerous endemic species. Based on our data, we recommend that Pemanggil Island should also be considered a protected area.

In the northwestern region, the Langkawi Archipelago (area 1; Fig. 2A) is designated as a Geopark under the United Nations Educational, Scientific, and Cultural Organization (UNESCO) and hence, has comprehensive sustainable development strategies in place (Norhayati et al. 2011). A small portion of area 2, namely Perlis State Park, is protected under category II but many of the indicator species in this area occur outside of the park especially in sensitive and relatively unprotected karst formations at Gua Kelam, Wang Kelian, Bukit Chabang, and Kuala Perlis. Similarly, the Selangor-Genting complex (area 7; Fig. 2A) contains numerous high-level PAs (categories 1a, 1b, and II), but the majority of these PAs are small, non-contiguous, and surrounded by urban development. Our results suggest that the amount and/or total area of PAs within these areas should be increased to encompass adjacent localities that have been shown to harbor sensitive species.

The remaining four areas (areas 3, 4, 5, and 10; Fig. 2A) are afforded relatively low levels of protection (category V–VI; Table 1) that allows continuous human interaction and sustainable use of natural resources. The most critical of these areas is Cameron Highlands in area 4, which has the highest number of EDGE species (Fig. 3A) and is among the most important area for tourism, agriculture, horticulture, and hydroelectric power generation in Peninsular Malaysia. Over the last decade, rapid and unregulated land conversion has led to a substantial loss of natural forests, resulting in severe environmental degradation characterized by surging cases of slope failure, floods, river pollution, and temperature increase (Rozimah & Khairulmaini 2016; Razali et al. 2018). Five critically endangered species of frogs and lizards occur in Cameron Highlands. One of those species, *Leptobrachella kecil* has the highest EDGE score (0.91) out of all assessed Peninsular Malaysian frogs and is only know of its type locality, which, has in recent years, been decimated and replaced by a condominium complex—this evolutionarily irreplaceable species is feared to be extinct. Other species endemic to this region are also known only from their type locality (i.e. *Kalophrynus yongi*, *Sphenomorphus cameronicus, Pseudocalotes rhaegal,* and *Tytthoscincus jaripendek*). Similar to Cameron Highlands, the southern Bintang range (area 3), Fraser’s Hill, and Genting Highlands also harbor numerous high-elevation and endemic species, many of which are threatened (Table 1). Although relatively less degraded compared to Cameron Highlands, these high-elevation areas are important tourist destinations due to the cool climate. It is therefore imperative that current and future development of these areas implement sufficient measures to protect the highly sensitive habitat and endemic fauna contained therein. Other areas, especially in the northwestern, eastern (states of Kelantan and Terengganu), and southeastern region (Seribuat Archipelago) were not only identified as areas of high conservation value (Figs. 1, 2) but also areas of biogeographic importance (Fig. 3; Table 2). Taken together, our results demonstrate north-south and east-west patterns of bioregionalization that are supported across numerous studies (Grismer & Pan 2008; Grismer et al. 2010b, 2015; Matsui et al. 2014; Chan et al. 2017, 2018). These patterns are largely modulated by the numerous mountain ranges that dominate the landscape of Peninsular Malaysia, forming barriers and corridors for gene flow we well as habitat refugia during periods of climatic fluctuations (Grismer et al. 2015; Chan et al. 2017; Chan & Brown 2020).

We note that the use of the third quartile threshold as a cut-off to determine high-value areas is subjective. However, it stands to reason that adjusting this threshold will only increase or decrease the number of retained areas without changing the metric value of each cell. This threshold is thus, analogous to a sensitivity parameter that can be adjusted according to the data. For our datasets, we found that the third quartile threshold retained sufficient areas for meaningful considerations. A second quartile threshold retained too many areas, while a top 10 percentile threshold retained too few. Another caveat is the inability of the EDGE analysis to analyze species that are categorized as data deficient (DD) in the IUCN Red List. Many DD species are newly described and are known to be highly endemic. Hence, their exclusion from analyses could potentially be detrimental to their survival. This underscores the urgent need to provide timely threat assessments for new species discoveries.

Our study demonstrates the utility of integrating spatial data and evolutionary history to identify areas for conservation priorities in Peninsular Malaysia. Successful conservation programs not only turn on the simple understanding of what species exist (i.e., taxonomy) and how they are distributed across a landscape (Mace 2004), but they are becoming more informed by robust evolutionary analyses. This enables researchers to identify species-rich landscapes and habitats with current and historically high rates of speciation and genetic diversity and target them for protection (Grismer et al. 2021). We identified Langkawi Geopark, the Benom-Krau complex, Endau-Rompin National Park, and Tioman Island as areas of high conservation value that are also afforded adequate levels of protection and therefore, need not be prioritized for urgent conservation action. Our data strongly indicate that Cameron Highlands and its surrounding regions should be given the highest conservation priority because of the high levels of range-restricted endemism, evolutionary distinct lineages, and threatened species. This could include the establishment of new PAs and/or the revision of existing PAs to a stricter category.

## Literature Cited

Aiken SR. 1994. Peninsular Malaysia’s Protected Areas’ Coverage, 1903–92: Creation, Rescission, Excision, and Intrusion. Environmental Conservation.

Barratt CD, Bwong BA, Onstein RE, Rosauer DF, Menegon M, Doggart N, Nagel P, Kissling WD, Loader SP. 2017. Environmental correlates of phylogenetic endemism in amphibians and the conservation of refugia in the Coastal Forests of Eastern Africa. Diversity and Distributions 23:875–887.

Carnaval AC et al. 2014. Prediction of phylogeographic endemism in an environmentally complex biome. Proceedings of the Royal Society B: Biological Sciences 281.

Chan KO, Abraham RK, Grismer JL, Grismer LL. 2018. Elevational size variation and two new species of torrent frogs from Peninsular Malaysia (Anura: Ranidae: *Amolops* Cope). Zootaxa 4434:250–264.

Chan KO, Ahmad N. 2009. Distribution and natural history notes on some poorly known frogs and snakes from Peninsular Malaysia. Herpetological Review 40:294–301.

Chan KO, Alexander AM, Grismer LL, Su Y-C, Grismer JL, Quah ESH, Brown RM. 2017. Species delimitation with gene flow: a methodological comparison and population genomics approach to elucidate cryptic species boundaries in Malaysian Torrent Frogs. Molecular Ecology 26:5435–5450.

Chan KO, Brown RM. 2020. Elucidating the drivers of genetic differentiation in Malaysian torrent frogs (Anura: Ranidae: Amolops): a landscape genomics approach. Zoological Journal of the Linnean Society 190:65–78.

Chan KO, Grismer LL. 2008. A new species of *Cnemaspis* Strauch 1887 (Squamata: Gekkonidae) from Selangor, Peninsular Malaysia. Zootaxa 57:49–58.

Chan KO, Grismer LL, Anuar S, Quah ESH, Muin MA, Savage AE, Grismer JL, Ahmad N, Remigio AC, Greer LF. 2010a. A new endemic rock gecko *Cnemaspis* Strauch 1887 (Squamata: Gekkonidae) from Gunung Jerai, Kedah, northwestern Peninsular Malaysia. Zootaxa 68:59–68.

Chan KO, Grismer LL, Sharma DS, Daicus B, Norhayati A. 2009. New herpetofaunal records for Perlis state park and adjacent areas. Malayan Nature Journal 61:277–284.

Chan KO, Muin MA, Anuar S, Andam J, Razak N, Aziz MA. 2019. First checklist on the amphibians and reptiles of Mount Korbu, the second highest peak in Peninsular Malaysia. Check List 15:1055–1069.

Chan KO, Muin MA, Badli-Sham BH, Fatihah-Syafiq M, Abraham RK, Ahmad A, Zakaria R. 2020. Identification and species delimitation of the enigmatic Marsh Frog *Pulchrana rawa* (Matsui, Mumpuni, and Hamidy, 2012): Second confirmed specimen and first country record for Malaysia. Journal of Herpetology 54:282–288.

Chan KO, van Rooijen J, Grismer LL, Belabut D, Muin MA, Jamaludin H, Gregory R, Norhayati A. 2010b. First report on the herpetofauna of Pulau Pangkor, Perak, Malaysia. Russian Journal of Herpetology 17:139–146.

Chan KO, Wood PLJ, Anuar S, Muin MA, Quah ESH, Sumarli AXY, Grismer LL. 2014. A new species of upland Stream Toad of the genus *Ansonia* Stoliczka, 1870 (Anura: Bufonidae) from northeastern Peninsular Malaysia. Zootaxa 3764:427–440.

Clements R, Ng PKL, Lu XX, Ambu S, Schilthuizen M, Bradshaw CJA. 2008. Using biogeographical patterns of endemic land snails to improve conservation planning for limestone karsts. Biological Conservation 141:2751–2764. Elsevier Ltd. Available from http://dx.doi.org/10.1016/j.biocon.2008.08.011.

Daru BH, Elliott TL, Park DS, Davies TJ. 2017. Understanding the Processes Underpinning Patterns of Phylogenetic Regionalization. Trends in Ecology and Evolution 32:845–860. Elsevier Ltd. Available from http://dx.doi.org/10.1016/j.tree.2017.08.013.

Daru BH, Karunarathne P, Schliep K. 2020. phyloregion: R package for biogeographical regionalization and macroecology. Methods in Ecology and Evolution 11:1483–1491.

Davis HR, Grismer LL, Klabacka RL, Muin MA, Quah ESH, Anuar S, Wood PLJ, Sites JW. 2016. The phylogenetic relationships of a new Stream Toad of the genus *Ansonia* Stoliczka, 1870 (Anura: Bufonidae) from a montane region in Peninsular Malaysia. Zootaxa 4103:137–153.

Edler D, Guedes T, Zizka A, Rosvall M, Antonelli A. 2017. Infomap bioregions: Interactive mapping of biogeographical regions from species distributions. Systematic Biology 66:197–204.

Evans LJ, Asner GP, Goossens B. 2018. Protected area management priorities crucial for the future of Bornean elephants. Biological Conservation 221:365–373. Elsevier. Available from https://doi.org/10.1016/j.biocon.2018.03.015.

Faith DP. 1992. Conservation evaluation and phylogenetic diversity. Biological Conservation 61:1–10.

FDPM. 2016. Forestry Department of Peninsular Malaysia Annual Report 2015. Kuala Lumpur.

Fenker J et al. 2020. Evolutionary history of Neotropical savannas geographically concentrates species, phylogenetic and functional diversity of lizards. Journal of Biogeography:1–13.

González-Orozco CE, Mishler BD, Miller JT, Laffan SW, Knerr N, Unmack P, Georges A, Thornhill AH, Rosauer DF, Gruber B. 2015. Assessing biodiversity and endemism using phylogenetic methods across multiple taxonomic groups. Ecology and Evolution 5:5177–5192.

Grismer LL et al. 2021. Phylogenetic partitioning of the third-largest vertebrate genus in the world, *Cyrtodactylus* Gray, 1827 (Reptilia; Squamata; Gekkonidae) and its relevance to taxonomy and conservation. Vertebrate Zoology 71:101–154.

Grismer LL, Anuar S, Quah ESH, Muin MA, Chan KO, Grismer JL, Ahmad N. 2010a. A new spiny, prehensile-tailed species of *Cyrtodactylus* (Squamata: Gekkonidae) from Peninsular Malaysia with a preliminary hypothesis of relationships based on morphology. Zootaxa 2625:40–52.

Grismer LL, Belabut DM, Quah ESH, Chan KO, Wood PL, Hasim R. 2014a. A new species of karst forest-adapted Bent-toed Gecko (genus *Cyrtodactylus* Gray, 1827) belonging to the *C. sworderi* complex from a threatened karst forest in Perak, Peninsular Malaysia. Zootaxa 3755:434–446.

Grismer LL, Chan KO. 2008. A new species of *Cnemaspis* Strauch 1887 (Squamata: Gekkonidae) from Pulau Perhentian Besar, Terengganu, Peninsular Malaysia. Zootaxa 1771:1–15.

Grismer LL, Chan KO, Grismer JL, Wood PL, Norhayati A, Ahmad N. 2010b. A checklist of the herpetofauna of the Banjaran Bintang, Peninsular Malaysia. Russian Journal of Herpetology 17:147–160.

Grismer LL, Chan KO, Quah ESH, Muin MA, Savage AE, Grismer JL, Ahmad N, Greer LF, Remegio AC. 2010c. Another new, diminutive Rock Gecko (*Cnemaspis* Strauch) from Peninsular Malaysia and a discussion of resource partitioning in sympatric species pairs. Zootaxa 66:55–66.

Grismer LL, Grismer JL, Wood PL, Ngo VT, Neang T, Chan KO. 2011a. Herpetology on the fringes of the Sunda shelf: a discussion of discovery, taxonomy, and biogeography. Bonner Zoologische Monographien 57:57–97.

Grismer LL, Pan KA. 2008. Diversity, endemism, and conservation of the amphibians and reptiles of southern Peninsular Malaysia and its offshore islands. Herpetological Review 39:270–281. Available from papers://1534bf8c-273e-4fe2-b9af-fcb670e73bd9/Paper/p2511.

Grismer LL, Quah ESH, Siler CD, Chan KO, Wood PL, Grismer JL, Anuar S, Ahmad N. 2011b. Peninsular Malaysia’s first limbless lizard: a new species of skink of the genus *Larutia* (Böhme) from Pulau Pinang with a phylogeny of the genus. Zootaxa 2799:29–40.

Grismer LL, Wood PL, Anuar S, Quah ESH, Muin MA, Chan KO, Sumarli AX, Loredo AI. 2015. Repeated evolution of sympatric, palaeoendemic species in closely related, co-distributed lineages of *Hemiphyllodactylus* Bleeker, 1860 (Squamata: Gekkonidae) across a sky-island archipelago in Peninsular Malaysia. Zoological Journal of the Linnean Society 174:859–876.

Grismer LL, Wood PL, Chan KO, Anuar S, Muin MA. 2014b. Cyrts in the city: a new Bent-toed gecko (genus *Cyrtodactylus*) is the only endemic species of vertebrate from Batu Caves, Selangor, Peninsular Malaysia. Zootaxa 3774:381–394.

Gumbs R, Gray CL, Böhm M, Hoffmann M, Grenyer R, Jetz W, Meiri S, Roll U, Owen NR, Rosindell J. 2020. Global priorities for conservation of reptilian phylogenetic diversity in the face of human impacts. Nature Communications 11:1–13. Springer US. Available from http://dx.doi.org/10.1038/s41467-020-16410-6.

Hashim Z, Abdullah SA, Nor SM. 2017. Stakeholders analysis on criteria for protected areas management categories in Peninsular Malaysia. IOP Conference Series: Earth and Environmental Science 91:012014.

Hoang DT, Chernomor O, von Haeseler A, Minh BQ, Le SV. 2017. UFBoot2: improving the ultrafast bootstrap approximation. Molecular Biology and Evolution 35:518–522. Available from http://academic.oup.com/mbe/article/doi/10.1093/molbev/msx281/4565479/UFBoot2-Improving-the-Ultrafast-Bootstrap.

Isaac NJB, Redding DW, Meredith HM, Safi K. 2012. Phylogenetically-Informed Priorities for Amphibian Conservation. PLoS ONE 7:1–8.

Isaac NJB, Turvey ST, Collen B, Waterman C, Baillie JEM. 2007. Mammals on the EDGE: Conservation priorities based on threat and phylogeny. PLoS ONE 2.

IUCN. 2019. The IUCN Red List of Threatened Species. Available from http://www.iucnredlist.org (accessed December 10, 2019).

Kalyaanamoorthy S, Minh BQ, Wong TKF, von Haeseler A, Jermiin LS. 2017. ModelFinder: fast model selection for accurate phylogenetic estimates. Nature Methods 14:587–589.

Katoh K, Standley DM. 2013. MAFFT multiple sequence alignment software version 7: Improvements in performance and usability. Molecular Biology and Evolution 30:772–780.

Kearse M et al. 2012. Geneious Basic: an integrated and extendable desktop software platform for the organization and analysis of sequence data. Bioinformatics 28:1647–9. Available from http://bioinformatics.oxfordjournals.org/content/28/12/1647.short.

Mace GM. 2004. The role of taxonomy in species conservation. Philosophical Transactions of the Royal Society B: Biological Sciences 359:711–719.

Mace GM, Gittleman JL, Purvis A. 2003. Preserving the Tree of Life. Science 300:1707–1709.

Matsui M. 2009. A new species of Kalophrynus with a unique male humeral spine from Peninsular Malaysia (Amphibia, Anura, Microhylidae). Zoological Science 26:579–585.

Matsui M, Belabut DM, Ahmad N. 2014. Two new species of fanged frogs from Peninsular Malaysia (Anura: Dicroglossidae). Zootaxa 3881:75–93.

Matsui M, Belabut DM, Ahmad N, Yong H-S. 2009. A new species of Leptolalax (Amphibia, Anura, Megophryidae) from Peninsular Malaysia. Zoological Science 26:243–247.

Miller M a, Pfeiffer W, Schwartz T. 2010. Creating the CIPRES Science Gateway for inference of large phylogenetic trees. 2010 Gateway Computing Environments Workshop, GCE 2010:1–8.

Minh BQ, Schmidt HA, Chernomor O, Schrempf D, Woodhams MD, Von Haeseler A, Lanfear R, Teeling E. 2020. IQ-TREE 2: New Models and Efficient Methods for Phylogenetic Inference in the Genomic Era. Molecular Biology and Evolution 37:1530–1534.

Mishler BD, Knerr N, González-Orozco CE, Thornhill AH, Laffan SW, Miller JT. 2014. Phylogenetic measures of biodiversity and neo- and paleo-endemism in Australian Acacia. Nature Communications 5:4473. Available from http://www.nature.com/doifinder/10.1038/ncomms5473.

Mittermeier R, Goettsch C, Robles Gil P. 2005. Megadiversity: Earth’s Biologically Wealthiest Nations1st Editio. CEMEX.

Myers N, Mittermeier RA, Mittermeier CG, da Fonseca GAB, Kent J. 2000. Biodiversity hotspots for conservation priorities. Nature 403:853–858.

Norhayati A, Chan KO, Daicus B, Samat A, Grismer LL, Mohd Izzuddin A. 2011. Potential biosites of significant importance in Langkawi Geopark: Terrestrial vertebrate fauna. Planning Malaysia 9:103–120.

NRE. 2016. Malaysia: National policy on biological diversity (2016–2025). Ministry of Natural Resources and Environment Malaysia (NRE), Putrajaya.

Pavoine S, Ollier S, Dufour AB. 2005. Is the originality of a species measurable? Ecology Letters 8:579–586.

Posadas P, Miranda Esquivel DR, Crisci J V. 2001. Using phylogenetic diversity measures to set priorities in conservation: An example from southern South America. Conservation Biology 15:1325–1334.

Quah ESH, Anuar S, Grismer LL, Wood. P. L. Jr., Azizah SMN. 2019a. Systematics and natural history of mountain reed snakes (genus *Macrocalamus*◻; Calamariinae). Zoological Journal of the Linnean Society:1–41.

Quah ESH, Anuar S, Grismer LL, Wood PL, Siti Azizah MNS, Muin MA. 2017. A new species of frog of the genus *Abavorana* Oliver, Prendini, Kraus & Raxworthy 2015 (Anura: Ranidae) from Gunung Jerai, Kedah, northwestern Peninsular Malaysia. Zootaxa 4320:272–288.

Quah ESH, Anuar SMS, Grismer LL, Muin MA, Chan KO, Grismer JL. 2011. Preliminary checklist of the herpetofauna of Jerejak Island, Penang, Malaysia. Malayan Nature Journal 63:595–600.

Quah ESH, Grismer LL, Wood. P. L. Jr., Thura MK, Oaks JR, Lin A. 2019b. Discovery of the westernmost population of the genus *Ansonia* Stoliczka (Anura, Bufonidae) with the description of a new species from the Shan Plateau of eastern Myanmar. Zootaxa 4656:545–571.

Ratnayeke S, Van Manen FT, Clements GR, Kulaimi NAM, Sharp SP. 2018. Carnivore hotspots in Peninsular Malaysia and their landscape attributes. PLoS ONE 13:e0194217.

Razali A, Syed Ismail SN, Awang S, Praveena SM, Zainal Abidin E. 2018. Land use change in highland area and its impact on river water quality: a review of case studies in Malaysia. Ecological Processes 7. Ecological Processes.

Redding DW, Mooers AO. 2006. Incorporating evolutionary measures into conservation prioritization. Conservation Biology 20:1670–1678.

Rosauer D, Laffan SW, Crisp MD, Donnellan SC, Cook LG. 2009. Phylogenetic endemism: A new approach for identifying geographical concentrations of evolutionary history. Molecular Ecology 18:4061–4072.

Rozimah R, Khairulmaini OS. 2016. Highland Regions – Land use Change Threat and Integrated River Basin Management. International Journal of Applied Environmental Sciences 11:1509–1521.

Sumarli AX, Grismer LL, Anuar S, Muin MA, Quah ESH. 2015. First report on the amphibians and reptiles of a remote mountain, Gunung Tebu in northeastern Peninsular Malaysia. Checklist 11:1–32.

Sumarli AX, Grismer LL, Wood PLJ, Ahmad AB, Rizal S, Ismail LH, Izam NAM, Ahmad N, Linkem CW. 2016. The first riparian skink (Genus: *Sphenomorphus* Strauch, 1887) from Peninsular Malaysia and its relationship to other Indochinese and Sundaic species. Zootaxa 4173:29.

Williams PH, Humphries CJ. 1994. Biodiversity, taxonomic relatedness, and endemism in conservation. Pages 269–287 in P. L. Forey, C. J. Humphries, and R. I. Vane-Wright, editors. Systematics and Conservation Evaluation. Oxford University Press, Oxford, UK.

